# Dose-Response Immunomodulatory Effects of Mesenchymal Stem Cells-Derived Culture-Conditioned Media in Acute Graft-versus-Host-Disease

**DOI:** 10.1101/2025.02.25.640019

**Authors:** Mohini Mendiratta, Meenakshi Mendiratta, Sujata Mohanty, Sandeep Rai, Ritu Gupta, Vatsla Dadhwal, Sameer Bakhshi, Deepam Pushpam, Mukul Aggarwal, Aditya Kumar Gupta, Ranjit Kumar Sahoo

## Abstract

**Background:** Mesenchymal stem cell-based therapy faces challenges that have driven interest in MSCs-derived culture-conditioned media (CCM) as a cell-free alternative. Our study aims to optimize the dose, and collection timing of CCM to enhance its therapeutic efficacy in aGVHD, while also standardizing co-culture conditions for CD3^+^ T-cell interaction with CCM.

**Material and Methods:** Human MSCs were isolated from BM and WJ and subsequently preconditioned under hypoxic conditions (1% O_2_) for 24 hours in a tri-gas incubator. Culture-conditioned media (CCM) was collected from both naive and hypoxia-preconditioned MSCs at 24, 48, and 72 hours and filtered using a 0.2 μm membrane filter. CD3+ T-cell were isolated from PBMNCs derived from aGVHD patients. These T-cell were co-cultured at varying densities (2*10^6^, 5*10^6^, and 10*10^6^ cells/ml) with different concentrations of CCM (25%, 50%, and 100%), and cell proliferation was assessed using the MTS assay. Furthermore, CD3+ T-cell proliferation and activation status were evaluated in a 2D co-culture model of CD3+ T-cell and CCM using flow cytometry.

**Results:** Our findings revealed that CCM collected at 48 hours, at a 50% concentration, exerted the most pronounced inhibitory effect on CD3+ T-cell proliferation, particularly at a density of 5*10^6^ cells/ml, irrespective of the MSCs source. Hypoxia preconditioning significantly enhanced the immunomodulatory effects, with WJ-MSCs^HYP^-CCM demonstrating superior efficacy in suppressing T-cell proliferation, increasing the CD4+/CD8+ T-cell ratio, and reducing CD4+ T-cell activation compared to BM-MSCs^HYP^-CCM.

**Conclusion:** These results emphasize the critical role of optimizing CCM collection timing and concentration to maximize therapeutic potential. Our study paves the way for the development of standardized, scalable, and effective cell-free therapies for aGVHD.

**Graphical Abstract:** 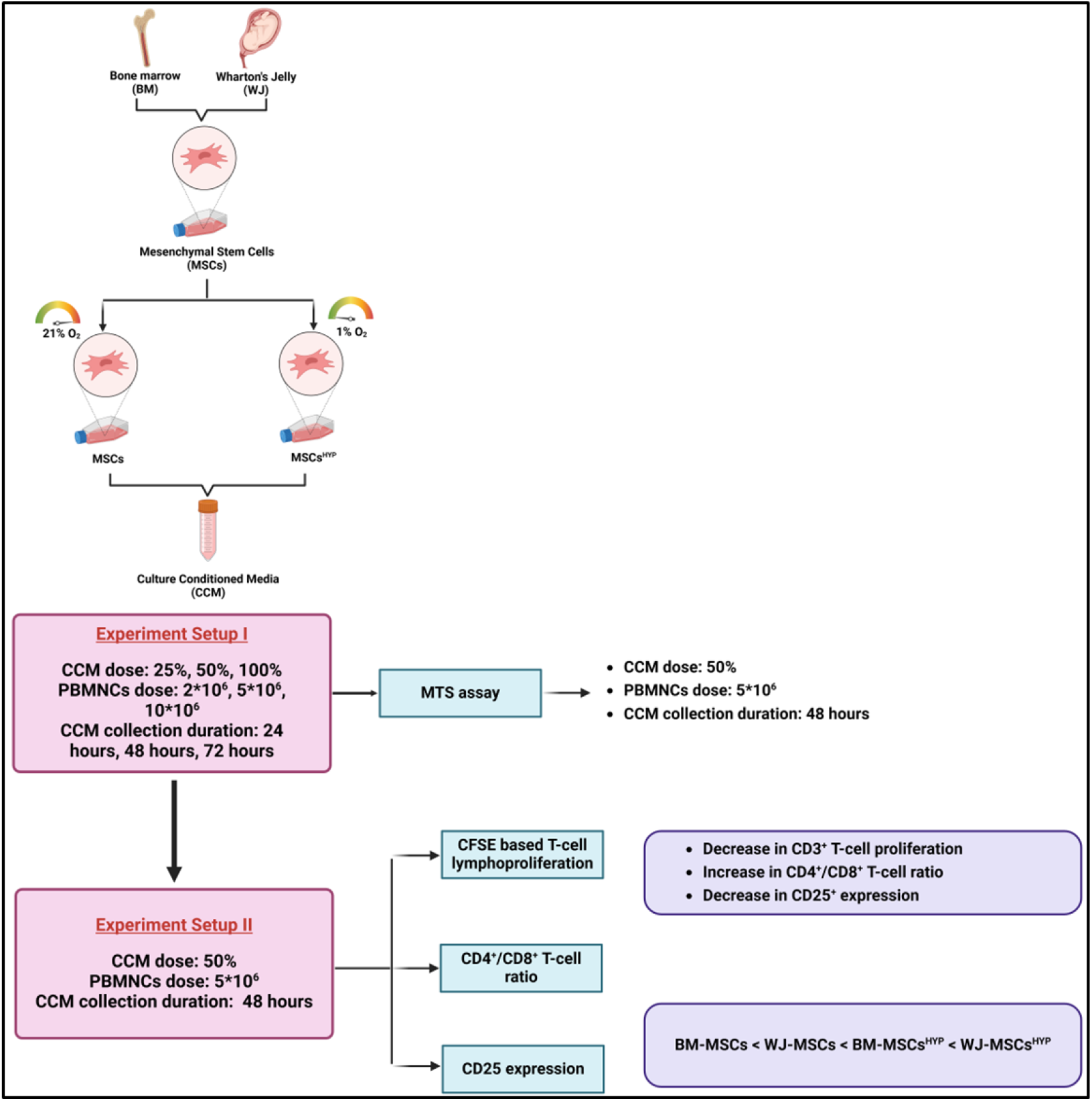

Optimum conditions for CCM dose, collection time point, and CD3^+^ T-cell dose for effective immunomodulatory effect of MSCs-derived CCM in aGVHD. (created using Biorender.com)

## Introduction

Mesenchymal stem cells have emerged as a promising therapeutic avenue due to their potent immunomodulatory properties (1). Although viable MSCs-based therapies show promise, challenges such as donor variability (2), cell viability (2,3), and inconsistent outcomes (3,4) have hindered their widespread application.

In recent years, the use of MSCs-derived culture-conditioned media (CCM) has emerged as a potential alternative, offering a cell-free therapeutic approach that circumvents the limitations while retaining their immunomodulatory potential. The therapeutic potential of CCM lies in its rich composition of bioactive molecules, including cytokines, chemokines, growth factors, and extracellular vesicles (EVs), which collectively contribute to immunomodulation (5) and tissue repair (6).

Previous studies indicated that the bioactivity of CCM can vary considerably based on factors such as the initial density of MSCs (7), culture duration (8), and the specific tissue source from which the MSCs are derived (9). Therefore, the dose-response relationship of CCM is particularly crucial, as insufficient doses may fail to achieve the desired therapeutic effect, while excessive doses could potentially disrupt immune homeostasis. A study demonstrated that a concentration of approximately 1-2*10^6^ MSCs per kilogram body weight is optimal for maximizing Tregs expansion and minimizing inflammatory responses (10).

Moreover, the optimization of the dose of CD3^+^ T-cell is equally essential for enhancing the therapeutic effects of MSCs-derived CCM. The interaction between CD3^+^ T-cell and MSCs-derived CCM establishes an immunological balance, and inappropriate dosing of PBMNCs may lead to insufficient regulatory control over the immune response or, conversely, excessive inflammatory reactions.

Additionally, the composition and potency of CCM are influenced by the timing of its collection, reflecting the dynamic secretory profile of MSCs under *in vitro* conditions (11,12). Studies have indicated that CCM harvested at specific intervals typically within 24 to 48 hours post-culture exhibits enhanced concentrations of key regulatory factors such as IL-6 and TGF-β, both of which are known to play significant roles in promoting anti-inflammatory pathways (13).

Our study aims to optimize the dose of MSCs-derived CCM and identify the appropriate time point for CCM collection, focusing on how these factors can be tailored to improve outcomes in patients suffering from aGVHD. Furthermore, our study seeks to standardize the conditions for direct 2D co-culture of CD3^+^ T-cell with

MSCs-derived CCM. By fine-tuning these parameters, we aim to provide a robust framework for developing MSCs-derived CCM as a standardized and effective therapeutic modality for aGVHD. This approach enhances the translational potential of MSC-derived therapies and aligns with the growing interest in cell-free solutions that are safer, scalable, and less prone to variability, offering new hope for improving outcomes in aGVHD management.

## Materials and Methods

### Collection of MSCs-derived CCM

Initially, human MSCs were isolated from bone marrow (BM) and Wharton’s Jelly (WJ) and preconditioned at 1% O_2_ for 24 hours, as described elsewhere (14). Before collecting the CCM from each group (MSCs, MSCs^HYP^), the FBS-containing culture medium was replaced with Stem Pro™ MSC SFM (Thermo Fisher Scientific, USA) to eliminate xenogenic components. After 24 hours, 48 hours, and 72 hours the CCM was collected and centrifuged at 3,500 rpm for 10 minutes and the supernatant was then filtered through a 0.2 μm syringe filter (Sigma-Aldrich, Saint Louis, MO, USA) to remove cell debris. The filtered CCM was then stored at -80°C for further experiments.

### Isolation and stimulation of CD3^+^ T-cell

Peripheral blood (PB) was collected from aGVHD patients in a sterile sodium heparin-coated vacutainer (Becton Dickinson, USA) and the peripheral blood mononuclear cells (PBMNCs) were isolated using Ficoll (Sigma-Aldrich, Saint Louis, MO, USA) density gradient centrifugation as described elsewhere. CD3^+^ T-cell was then isolated from PBMNCs by negative selection using a Pan T cell isolation kit, human (Miltenyi Biotec, USA), following the manufacturer’s instructions. Subsequently, CD3^+^ T-cell was stimulated with 1.0 μg/ml of Phytohemagglutinin (PHA) (Sigma-Aldrich, Saint Louis, MO, USA) and 50 IU/ml of IL-2 (Thermo Fisher Scientific, USA) in 1X Rosewell Park’s Memorial Institute (RPMI)-1640 medium supplemented with L-glutamine (Thermo Fisher Scientific, USA), 10% FBS (Thermo Fisher Scientific, USA), 1% antibiotic-antimycotic solution (Thermo Fisher Scientific, USA) at 37°C, 5% CO_2_ for 48 hours (15).

### MTS assay for T-cell proliferation

The proliferation of CD3^+^ T-cell was assessed using the 3- (4,5-dimethylthiazol-2-yl)-5-(3-carboxymethoxyphenyl)-2-(4-sulfophenyl)-2H-tetrazolium (MTS) (Abcam, USA) following the manufacturer’s protocol. Briefly, 10μl of MTS (Abcam, USA) solution was added directly to each well of 96-well plates after 72 hours of co-culture, containing PHA-stimulated and non-activated (control) cells, along with CCM collected from MSCs, MSCs^HYP^ at different time points-24, 48, and 72 hours. The plates were then incubated for 3 hours at 37°C. Following incubation, absorbance was measured at 490 nm using an ELISA reader (Bio-Rad, USA), with the blank consisting of culture medium alone. All experiments were conducted with 5 healthy individuals and 5 aGVHD patients, resulting in 5 biological replicates of each. Within each biological replicate, experiments were conducted in triplicates (15).

### CFSE T-cell proliferation assay

CD3^+^ T-cell were labeled with 1 µM Cell Trace^TM^ CFSE dye (Becton Dickinson, USA) for 20 minutes at 37°C and followed by activation with PHA (1 µg/ml) (Sigma-Aldrich, Saint Louis, MO, USA) and IL-2 (50 IU/ml) (Sigma-Aldrich, Saint Louis, MO, USA) in 1X RPMI-1640 complete medium supplemented with L-glutamine (Thermo Fisher Scientific, USA), at 37°C, 5% CO_2_ for 48 hours. CFSE-labelled activated T-cell were treated with CCM (MSCs, MSCs^HYP^) in a 1:1 ratio with 1X RPMI-1640 complete medium (Thermo Fisher Scientific, USA) for 3 days.

After 3 days of co-culture, the proliferation of T-cell was assessed using a DxFlex flow cytometer (Beckman Coulter, USA), and the data was analyzed using Kaluza software version 2.1 (Beckman Coulter, USA). CFSE-labelled activated T-cell in the absence of CCM were used as a control for the normalization of the proliferation of T-cell in the direct co-culture. All experiments were conducted with 5 biological replicates of each aGVHD cohort (15).

### Statistical analysis

All statistical analyses were conducted using GraphPad Prism version 8.4.3. One-way and Tukey’s post hoc tests were used to compare three or more groups. Data was shown as Mean±S.D. and a p-value of ≤ 0.05 was considered statistically significant.

## Results

### 48 hours CCM at 50% concentration achieves maximum inhibition of CD3^+^ T-cell proliferation at 5 × 10^6^ cells/ml

In this study, we aimed to determine the optimal dose of MSCs-derived CCM (BM-MSCs-CCM, BM-MSCs^HYP^-CCM, WJ-MSCs-CCM, and WJ-MSCs^HYP^-CCM) (25-100% CCM) and aGVHD patients-derived CD3^+^ T-cell (2-10*10^6^ cells), the differential immunomodulatory effects of CCM collected at varying time points (24-72 hours) by assessing the cell proliferation in a 2D cellular model of aGVHD using MTS assay.

Our findings revealed that both BM-MSCs-CCM and BM-MSCs^HYP^-CCM collected at 24 hours did not significantly reduce CD3^+^ T-cell proliferation, regardless of the CCM and CD3^+^ T-cell doses. In contrast, CCM collected at 48 hours demonstrated a notable reduction in cell proliferation at 25% and 50% CCM concentrations across varying CD3^+^ T-cell densities (2–10 × 10^6^ cells/ml), with 50% CCM exhibiting greater inhibitory potency compared to 25% CCM. The most pronounced reduction in proliferation was observed at 5 × 10^6^ cells/ml compared to 2 × 10^6^ cells/ml in both BM-MSCs-CCM (0.1118 vs. 0.1164; *p* ≤ 0.05) (Figure 1A-C) and BM-MSCs^HYP^-CCM (0.0884 vs. 0.113; p ≤0.0001) (Figure 2A-C), followed by a plateau effect at 10 × 10^6^ cells/ml (0.1118 vs. 0.1118; *p* > 0.999), (0.0884 vs 0.0884; p > 0.999) when treated with 50% CCM. Interestingly, 100% CCM did not result in a significant reduction in proliferation at any CD3^+^ T-cell concentration. Moreover, both CCM collected at 72 hours did not exhibit a significant inhibitory effect on CD3^+^ T-cell proliferation, irrespective of the CCM or CD3^+^ T-cell dose.

**Figure 1.**
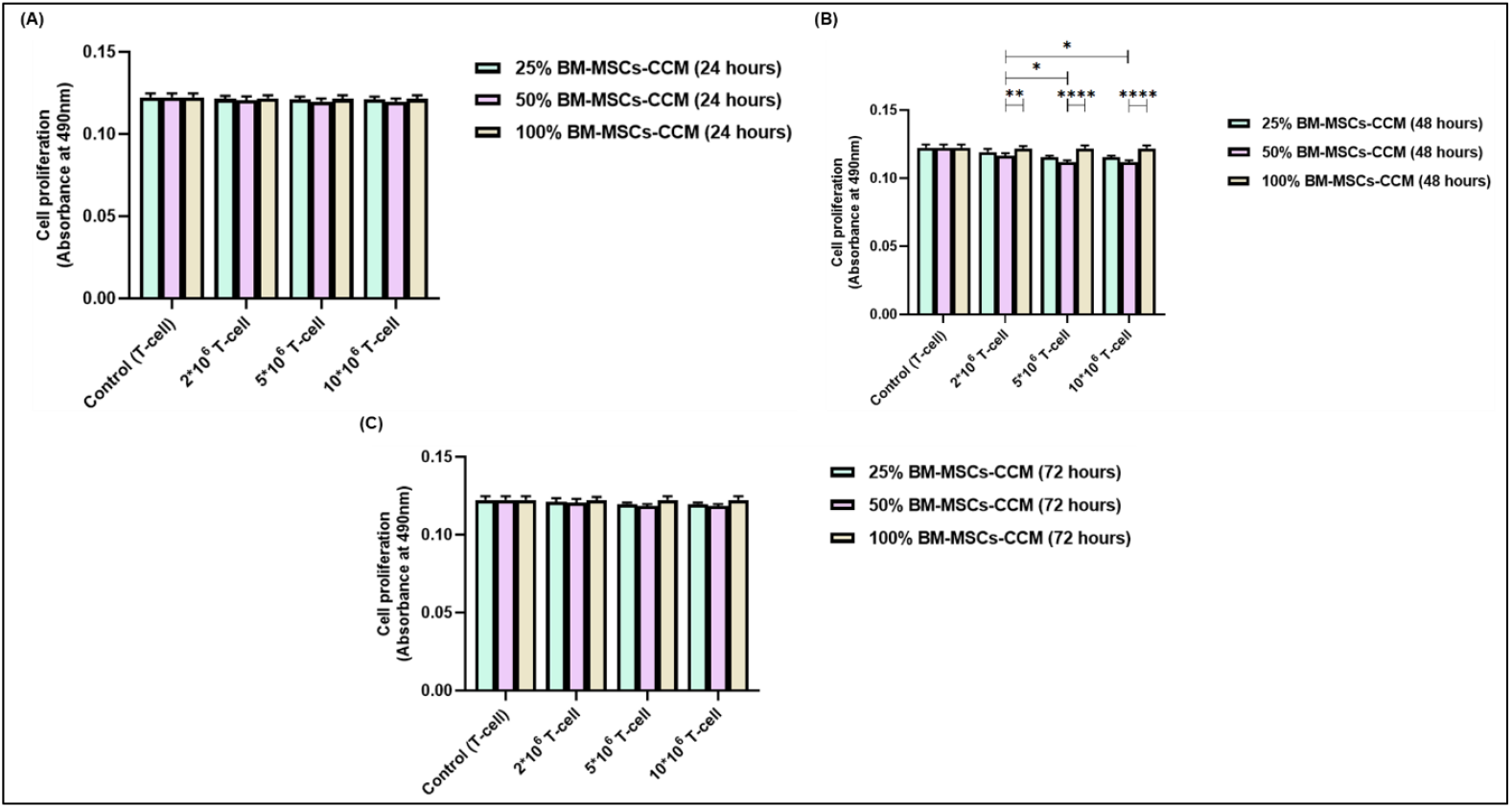
Proliferation of aGVHD patients derived CD3^+^ T-cell with and without different doses of BM-MSCs-CCM (25%, 50%, 100%) at various doses of CD3^+^ T-cell (2*10^6^, 5*10^6^, 10*10^6^ cells) at different time-points using MTS assay. The bar graph shows their proliferation at (A) 24 hours. (B) 48 hours. (C) 72 hours. Data shown represent the Mean±S.D of 5 independent experiments performed with T-cell derived from 5 different donors (biological replicates), with each experiment conducted in triplicates (technical replicates). Statistical analysis: Tukey’s multiple comparisons test; *≤0.05; ****≤0.0001. *Abbreviations: CCM: Culture-conditioned media*

**Figure 2.**
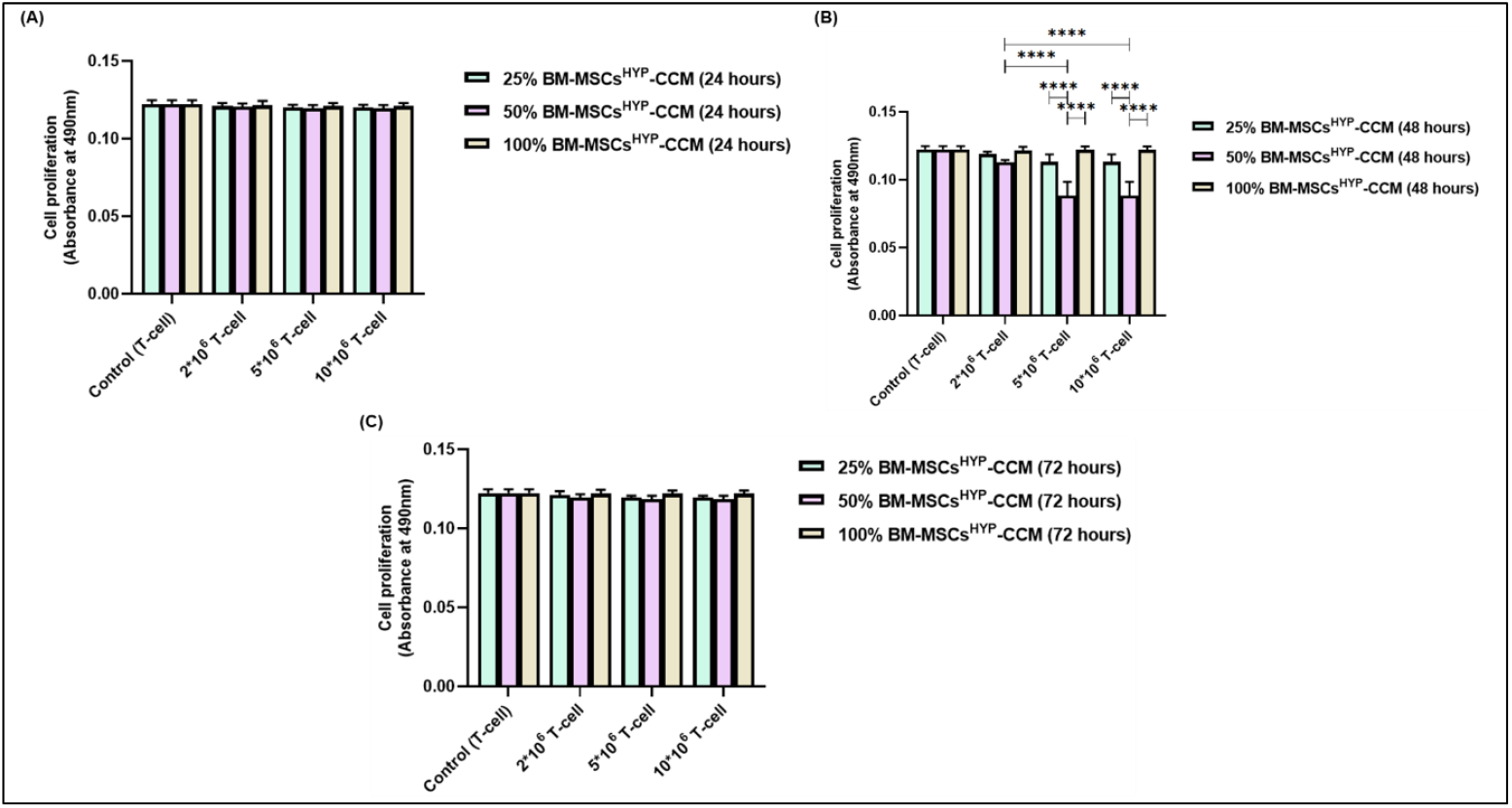
Proliferation of aGVHD patients derived CD3^+^ T-cell with and without different doses of BM-MSCs^HYP^-CCM (25%, 50%, 100%) at various doses of CD3^+^ T-cell (2*10^6^, 5*10^6^, 10*10^6^ cells) at different time-points using MTS assay. The bar graph shows their proliferation at (A) 24 hours. (B) 48 hours. (C) 72 hours. Data shown represent the Mean±S.D of 5 independent experiments performed with T-cell derived from 5 different donors (biological replicates), with each experiment conducted in triplicates (technical replicates). Statistical analysis: Tukey’s multiple comparisons test; ****≤0.0001. *Abbreviations: CCM: Culture-conditioned media*

Both WJ-MSCs-CCM and WJ-MSCs^HYP^-CCM collected at 24 hours did not exhibit a significant reduction in CD3^+^ T-cell proliferation across all tested conditions. In contrast, CCM collected at 48 hours significantly reduced cell proliferation at 25% and 50% concentrations across various CD3^+^ T-cell densities, whereas 100% CCM showed no significant inhibitory effect at any CD3^+^ T-cell concentration. Similarly, CCM collected at 72 hours did not result in a significant decrease in cell proliferation, regardless of the dose of CCM or CD3^+^ T-cell.

In both WJ-MSCs-CCM and WJ-MSCs^HYP^-CCM, the 50% CCM concentration was consistently more effective than 25% CCM in reducing cell proliferation across all CD3^+^ T-cell densities. The most pronounced inhibitory effect of 50% CCM was observed at 5 × 10^6^ CD3^+^ T-cell/ml compared to 2 × 10^6^ cells/ml in WJ-MSCs-CCM (0.100 vs. 0.115; *p* ≤ 0.0001) (Figure 3A-C) and WJ-MSCs^HYP^-CCM (0.0672 vs. 0.1052; *p* ≤ 0.0001) (Figure 4A-C). This effect reached a plateau at 10 × 10^6^ cells/ml (0.100 vs. 0.100; *p* > 0.999 for WJ-MSCs-CCM; 0.0672 vs. 0.0672; *p* > 0.999 for WJ-MSCs^HYP^-CCM). The key findings of optimization of dose of CCM, collection timing, and aPBMNCs dose is summarized in Table 1.

**Figure 3.**
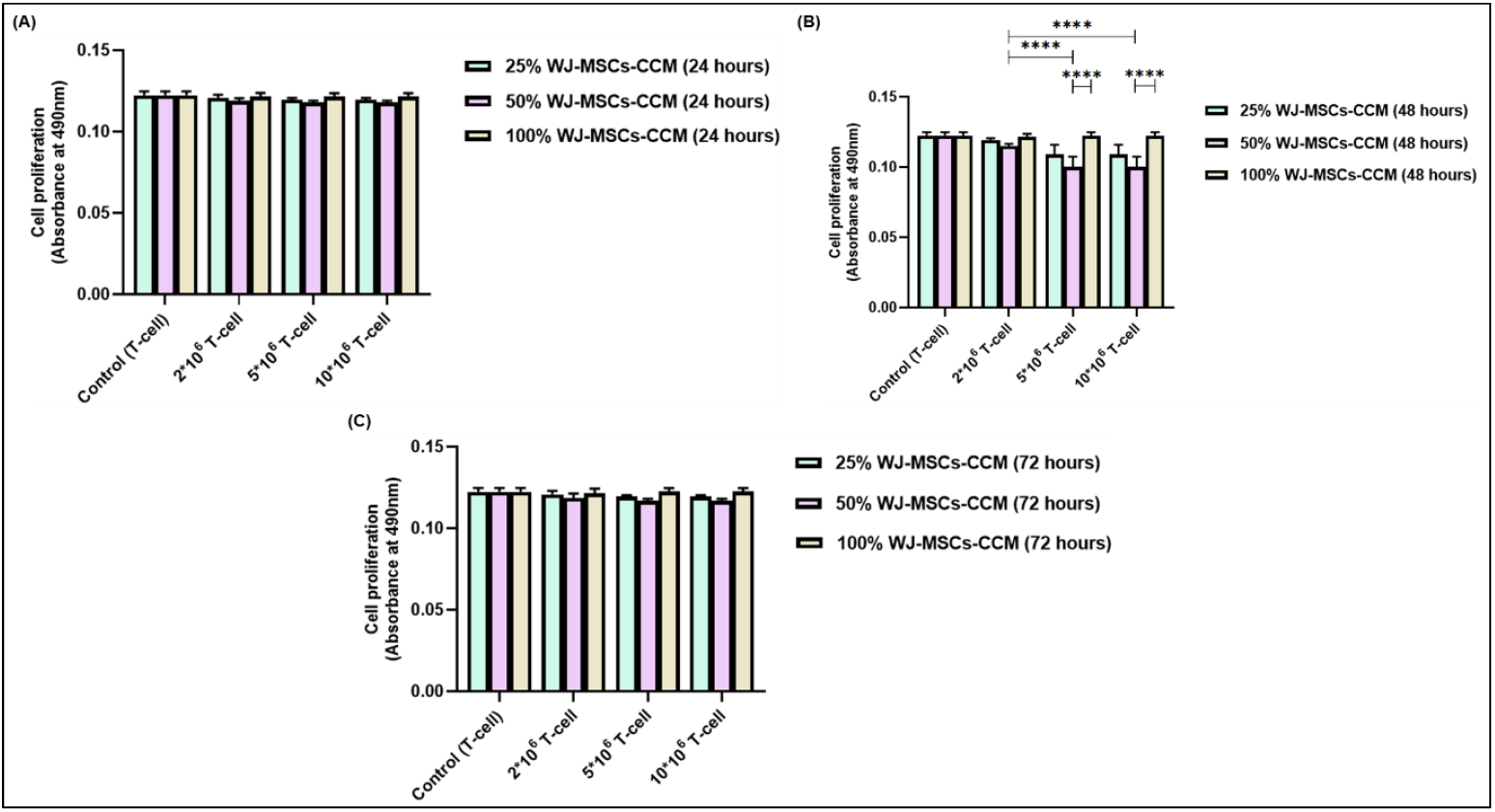
Proliferation of aGVHD patients derived CD3^+^ T-cell with and without different doses of WJ-MSCs-CCM (25%, 50%, 100%) at various doses of CD3^+^ T-cell (2*10^6^, 5*10^6^, 10*10^6^ cells) at different time-points using MTS assay. The bar graph shows their proliferation at (A) 24 hours. (B) 48 hours. (C) 72 hours. Data shown represent the Mean±S.D of 5 independent experiments performed with T-cell derived from 5 different donors (biological replicates), with each experiment conducted in triplicates (technical replicates). Statistical analysis: Tukey’s multiple comparisons test; ****≤0.0001. *Abbreviations: CCM: Culture-conditioned media*

**Figure 4.**
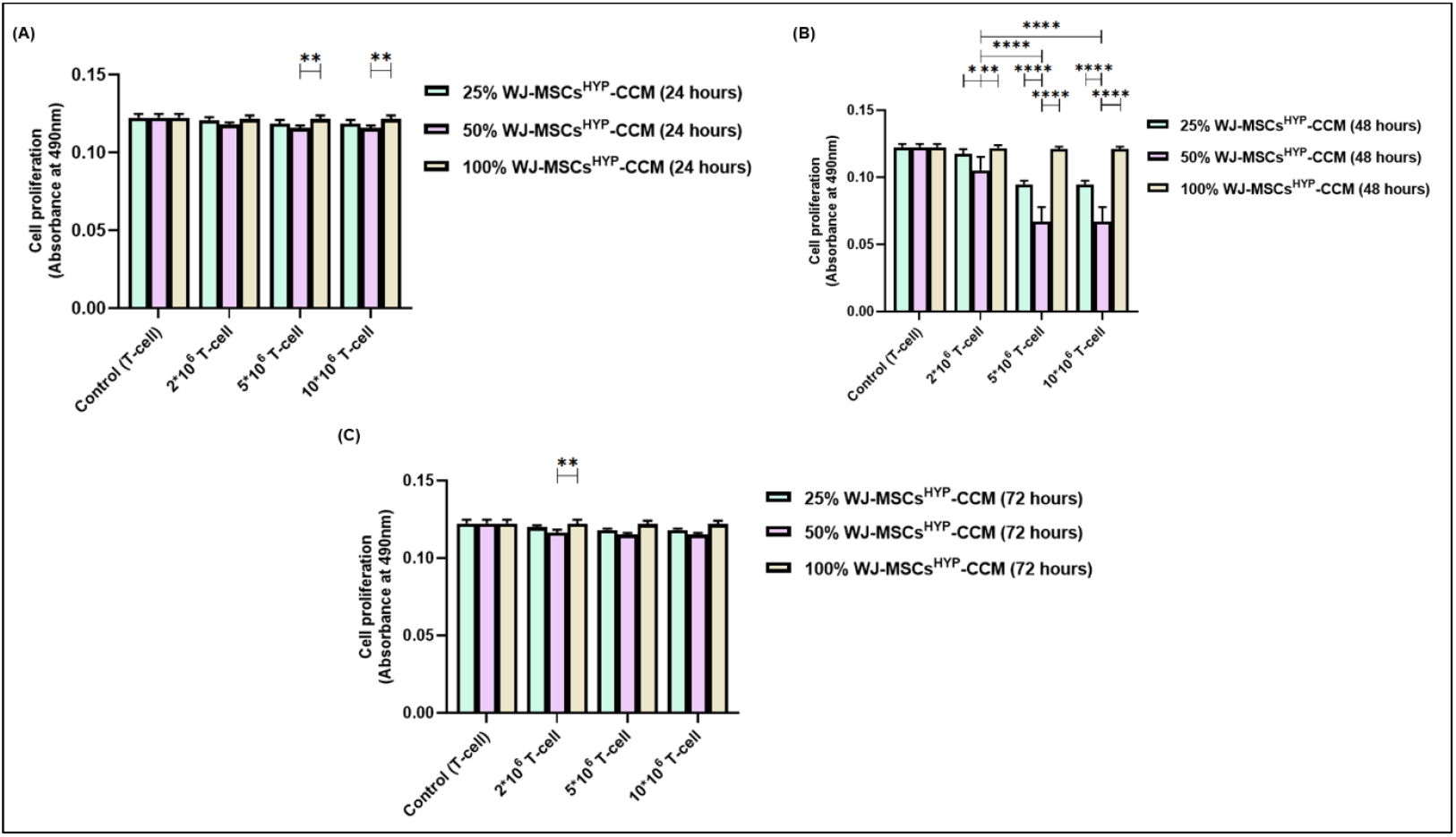
Proliferation of aGVHD patients derived CD3^+^ T-cell with and without different doses of WJ-MSCs^HYP^-CCM (25%, 50%, 100%) at various doses of CD3^+^ T-cell (2*10^6^, 5*10^6^, 10*10^6^ cells) at different time-points using MTS assay. The bar graph shows their proliferation at (A) 24 hours. (B) 48 hours. (C) 72 hours. Data shown represent the Mean±S.D of 5 independent experiments performed with T-cell derived from 5 different donors (biological replicates), with each experiment conducted in triplicates (technical replicates). Statistical analysis: Tukey’s multiple comparisons test; *≤0.05; **≤0.01; ****≤0.0001. *Abbreviations: CCM: Culture-conditioned media*

**Table 1:** Summary of key findings on optimization of CCM (dose and collection time points) and CD3^+^ T-cell dose for immunomodulation in aGVHD.

| Characteristics | Inference |
| --- | --- |
| CCM dose | 50% |
| CCM collection time point | 48 hours |
| CD3<sup>+</sup> T-cell dose | 5*10<sup>6</sup> cells/ml |

### WJ-MSCs^HYP^-CCM outperforms in inhibition of T-cell proliferation and activation

Our study demonstrated that MSCs-derived CCM collected at 48 hours with a 50% concentration exerted the most pronounced inhibitory effect on the proliferation of CD3^+^ T-cell (5 × 10^6^ cells), irrespective of the MSCs source. Flow cytometric analysis further revealed that both BM-MSCs-CCM and WJ-MSCs-CCM significantly suppressed CD3^+^ T-cell proliferation, with WJ-MSCs-CCM showing greater efficacy compared to BM-MSCs-CCM (51.246% vs. 57.33%; p ≤ 0.0001). Hypoxia preconditioning further enhanced the inhibitory effect of both MSCs sources, with WJ-MSCs^HYP^-CCM demonstrating superior suppression compared to BM-MSCs^HYP^-CCM (39.186% vs. 46.604%; p ≤ 0.0001) (Figure 5A).

**Figure 5.**
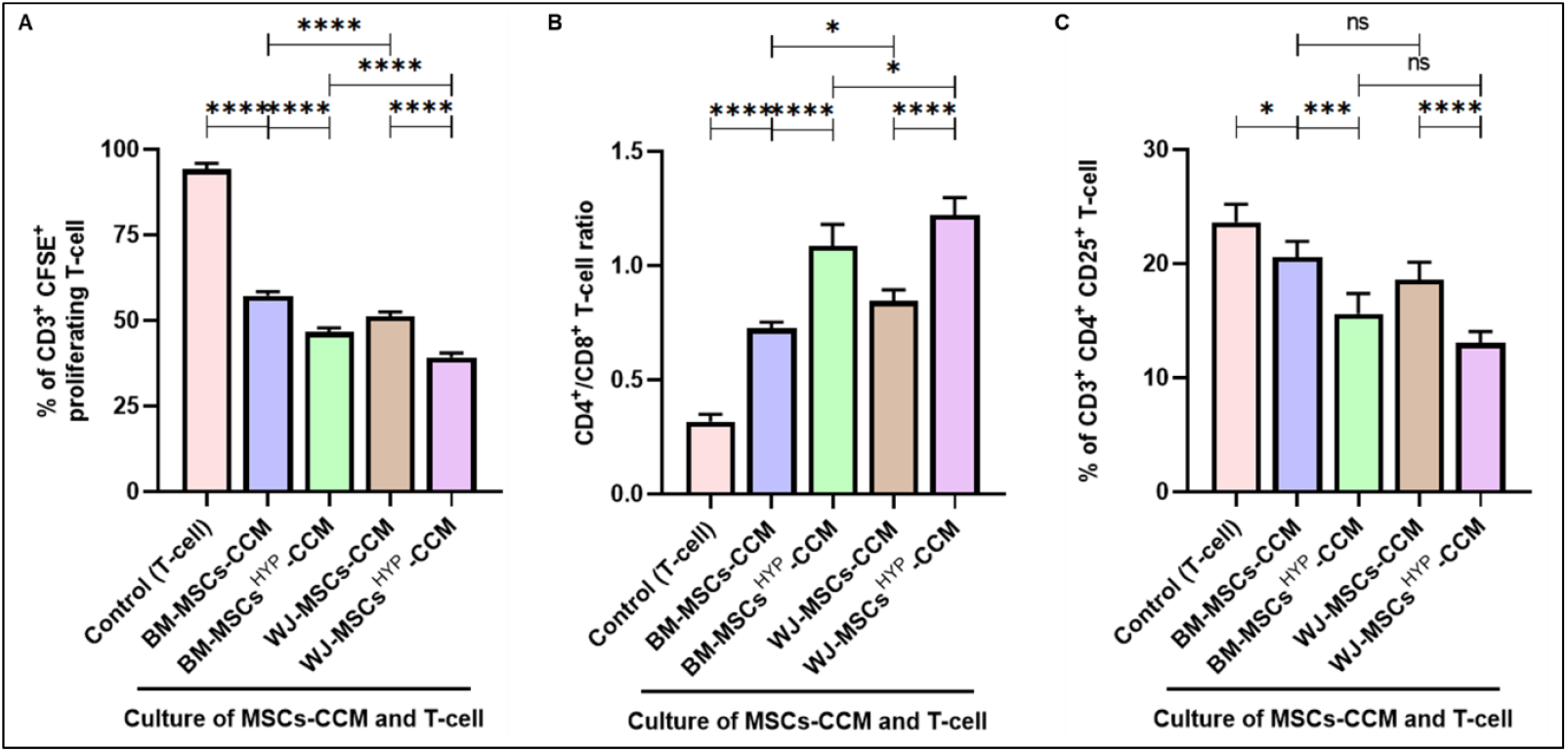
Proliferation of CD3+ T-cell (5 × 10^6^ cells/ml) derived from aGVHD patients after 72 hours of culture with 50% CCM collected at 48 hours. The bar graphs depict (A) % of CFSE+ proliferating CD3+ T-cell (B) CD4+/CD8+ T-cell ratio, and (C) % of CD3+ CD4+ CD25+ T-cell. Data shown represent the Mean±S.D of 5 independent experiments performed with T-cell derived from 5 different donors (biological replicates), with each experiment conducted in triplicates (technical replicates). Statistical analysis: Tukey’s multiple comparisons test; *≤0.05; ****≤0.0001. *Abbreviations: CCM: Culture-conditioned media*

The CD4^+^/CD8^+^ T-cell ratio was notably increased following treatment with CCM, with WJ-MSCs-CCM showing a more pronounced effect than BM-MSCs-CCM (0.845 vs. 0.726; p ≤ 0.05). Hypoxia preconditioning further amplified this effect, with WJ-MSCs^HYP^-CCM significantly outperforming BM-MSCs^HYP^-CCM (1.221 vs. 1.085; p ≤ 0.05) (Figure 5B). These results indicate that WJ-MSCs-CCM, particularly under hypoxic conditions, is highly effective in modulating the balance between CD4^+^ and CD8^+^ T-cell populations.

Additionally, CCM alleviated the activation status of CD4^+^ T-cell, with a stronger inhibitory effect observed for WJ-MSCs^HYP^-CCM compared to BM-MSCs^HYP^-CCM (13.056% vs. 18.61%; p ≤ 0.0001). Similar trends were observed for BM-MSCs^HYP^-CCM (18.61% vs. 20.584%; p ≤ 0.0001). However, no significant difference was noted between the naïve WJ-MSCs-CCM and BM-MSCs-CCM in terms of their effects on CD4^+^ T-cell activation (Figure 5C).

These findings highlight the enhanced immunomodulatory potential of hypoxia-preconditioned MSCs-derived CCM, especially from WJ-MSCs, across key immune parameters such as T-cell proliferation, CD4^+^/CD8^+^ T-cell ratio, and CD4^+^ T-cell activation. This underscores the therapeutic potential of WJ-MSCs^HYP^-CCM as a promising immunomodulatory agent for managing immune dysregulation in inflammatory conditions.

## Discussion

Our study highlights the immunomodulatory potential of MSCs-derived CCM in modulating CD3^+^ T-cell proliferation in aGVHD. This aligns with previous research that has consistently demonstrated the potent immunosuppressive effects of MSCs (16) and their secreted factors (17) on T-cell populations, highlighting their potential therapeutic applications in inflammatory conditions and transplant settings (18). The findings demonstrated that CCM collected at 48 hours with a 50% concentration exerts the most significant inhibition of CD3^+^ T-cell proliferation, particularly at 5*10^6^ cells/ml, irrespective of the MSCs source. These align with previous studies emphasizing the importance of optimizing MSCs derived-secretome components (19,20), collection time points (18,19), and dosage (19,21) to achieve maximum therapeutic efficacy. Additionally, the plateau effect observed at higher T-cell densities (10 × 10^6^ cells/ml) indicates that the immunosuppressive capacity of CCM is dose-dependent and may be limited by the saturation of receptor-mediated pathways.

Our study underscores the importance of the CCM collection’s concentration and timing. Notably, CCM collected at 24 hours did not yield significant inhibitory effects on T-cell proliferation, which is consistent with findings that suggest MSCs require a certain maturation period to secrete effective immunomodulatory factors (22). The mechanisms through which MSCs exert their immunomodulatory effects involve various soluble factors and cell contact-dependent interactions. It has been established that factors such as PGE2 and IDO play critical roles in mediating these effects, but their secretion may be contingent upon the activation state of the MSCs over time (22,23).

Our results further demonstrate that while 100% CCM failed to inhibit T-cell proliferation, a 50% concentration was significantly more effective than lower concentrations, reinforcing the idea that an optimal range exists for MSCs-derived factors to exert their immunosuppressive effects. This aligns with earlier findings that MSCs secrete dynamic profiles of cytokines, chemokines, and growth factors depending on the culture conditions and time points (24) and due to the presence of high concentrations of factors that potentially induce feedback inhibition or alter the balance of bioactive components (25). This is particularly relevant for clinical applications where precise dosing may enhance therapeutic outcomes.

The comparative analysis between BM-MSCs and WJ-MSCs revealed that WJ-MSCs-CCM significantly reduced T-cell proliferation, with hypoxia preconditioning further amplifying their inhibitory effects. This observation is consistent with reports that WJ-MSCs possess a more robust immunomodulatory profile due to their primitive nature and higher secretion of anti-inflammatory cytokines and growth factors (26,27). Furthermore, WJ-MSCs^HYP^ had an enhanced ability to suppress T-cell proliferation and reduce T-cell activation. This aligns with studies showing that hypoxic conditions upregulate HIF-1α-dependent pathways, enhancing the production of immunosuppressive molecules such as IDO, IL-10, TGF-β, and PGE2 (28–30).

The increased CD4^+^/CD8^+^ T-cell ratio observed with WJ-MSCsHYP-CCM treatment further underscores its potential to restore immune balance in aGVHD. This shift in T-cell populations may facilitate the generation of Tregs, a key mechanism for reducing effector T-cell activity and promoting immune tolerance (17). Additionally, WJ-MSCs^HYP^-CCM reduces CD25 expression, highlighting its role in attenuating the hyperactivation of effector T cells, a hallmark of aGVHD pathogenesis (31).

In summary, our study highlights the tailored application of MSCs-derived CCM in the context of managing immune responses in aGVHD. The significant immunomodulatory effects observed at the specified concentrations and timings illuminate the intricate dynamics governing T-cell biology and propose that further elucidation of the underlying mechanisms could enhance therapeutic strategies. The modulation of key immune parameters such as T-cell activation and ratios presents an opportunity for MSC-derived therapies to mitigate adverse immune responses and promote a balanced immune environment conducive to recovery and healing.

## Conclusion

In conclusion, our study reinforces the potential of MSCs-derived CCM as an innovative approach to immunotherapy. The optimization of CCM collection parameters and understanding of the differential effects based on MSCs sources provide critical insights for future research aimed at harnessing the therapeutic benefits of MSCs in clinical practice.

## Author’s Contributions

MM^1^ performed the experiments, acquired and analyzed data, interpreted the results, and wrote the manuscript. MM^2^ performed the experiments and acquired and analyzed data. SR and RG contributed to data interpretation and analysis. SM provided resources, conceptualized the study, and designed and supervised the experiments. VD, SB, DP, MA, and AKG provided patient samples and their clinical details. RKS provided funding and resources, conceptualized the study, designed and supervised the experiments, analyzed data, and edited the manuscript. All authors critically reviewed and approved the final version of the manuscript.

## Data availability

The data that support the findings of this study will be made available from the corresponding author upon reasonable request.

## Acknowledgment

The authors express their gratitude to the All India Institute of Medical Sciences (AIIMS), New Delhi, India for facilitating the execution of the study. A graphical abstract was created using Biorender.com.

## Abbreviations

MSCs: Mesenchymal Stem Cells
BM: Bone Marrow
WJ: Wharton’s Jelly
CCM: Culture-Conditioned Media
FBS: Fetal Bovine Serum
CO_2_: Carbon dioxide
PB: Peripheral Blood
PBMNCs: Peripheral Blood Mononuclear Cells
PHA: Phytohemagglutinin
IL-2: Interleukin-2
aGVHD: Acute Graft-versus-Host-Disease
MTS: 3-(4,5-dimethylthiazol-2-yl)-5-(3-carboxymethoxyphenyl)-2-(4-sulfophenyl)-2H-tetrazolium
CFSE: Carboxyfluorescein Succinimidyl Ester

## References

1. Müller L, Tunger A, Wobus M, von Bonin M, Towers R, Bornhäuser M, et al. Immunomodulatory Properties of Mesenchymal Stromal Cells: An Update. Front Cell Dev Biol. 2021;9:637725.

2. Baldari S, Di Rocco G, Piccoli M, Pozzobon M, Muraca M, Toietta G. Challenges and Strategies for Improving the Regenerative Effects of Mesenchymal Stromal Cell-Based Therapies. Int J Mol Sci. 2017 Oct 2;18(10):2087.

3. Mei R, Wan Z, Yang C, Shen X, Wang R, Zhang H, et al. Advances and clinical challenges of mesenchymal stem cell therapy. Front Immunol [Internet]. 2024 Jul 19 [cited 2025 Jan 25];15. Available from: https://www.frontiersin.org/journals/immunology/articles/10.3389/fimmu.2024.1421854/full

4. Shan Y, Zhang M, Tao E, Wang J, Wei N, Lu Y, et al. Pharmacokinetic characteristics of mesenchymal stem cells in translational challenges. Sig Transduct Target Ther. 2024 Sep 13;9(1):1–27.

5. Peshkova M, Korneev A, Suleimanov S, Vlasova II, Svistunov A, Kosheleva N, et al. MSCs’ conditioned media cytokine and growth factor profiles and their impact on macrophage polarization. Stem Cell Research & Therapy. 2023 May 25;14(1):142.

6. Gunawardena TNA, Rahman MT, Abdullah BJJ, Abu Kasim NH. Conditioned media derived from mesenchymal stem cell cultures: The next generation for regenerative medicine. J Tissue Eng Regen Med. 2019 Apr;13(4):569–86.

7. Pawitan JA. Prospect of Stem Cell Conditioned Medium in Regenerative Medicine. BioMed Research International. 2014;2014(1):965849.

8. Bogatcheva NV, Coleman ME. Conditioned Medium of Mesenchymal Stromal Cells: A New Class of Therapeutics. Biochemistry (Mosc). 2019 Nov;84(11):1375– 89.

9. Whittaker TE, Nagelkerke A, Nele V, Kauscher U, Stevens MM. Experimental artefacts can lead to misattribution of bioactivity from soluble mesenchymal stem cell paracrine factors to extracellular vesicles. Journal of Extracellular Vesicles. 2020;9(1):1807674.

10. Bonig H, Verbeek M, Herhaus P, Braitsch K, Beutel G, Schmid C, et al. Real-world data suggest effectiveness of the allogeneic mesenchymal stromal cells preparation MSC-FFM in ruxolitinib-refractory acute graft-versus-host disease. Journal of Translational Medicine. 2023 Nov 21;21:837.

11. Acuto S, Lo Iacono M, Baiamonte E, Lo Re R, Maggio A, Cavalieri V. An optimized procedure for preparation of conditioned medium from Wharton’s jelly mesenchymal stromal cells isolated from umbilical cord. Front Mol Biosci. 2023 Oct 2;10:1273814.

12. Montero-Vilchez T, Sierra-Sánchez Á, Sanchez-Diaz M, Quiñones-Vico MI, Sanabria-de-la-Torre R, Martinez-Lopez A, et al. Mesenchymal Stromal Cell-Conditioned Medium for Skin Diseases: A Systematic Review. Front Cell Dev Biol [Internet]. 2021 Jul 23 [cited 2025 Jan 25];9. Available from: https://www.frontiersin.org/journals/cell-and-developmental-biology/articles/10.3389/fcell.2021.654210/full

13. Kay AG, Long G, Tyler G, Stefan A, Broadfoot SJ, Piccinini AM, et al. Mesenchymal Stem Cell-Conditioned Medium Reduces Disease Severity and Immune Responses in Inflammatory Arthritis. Sci Rep. 2017 Dec 21;7(1):18019.

14. Mendiratta M, Mendiratta M, Ganguly S, Rai S, Gupta R, Kumar L, et al. Concurrent hypoxia and apoptosis imparts immune programming potential in mesenchymal stem cells: Lesson from acute graft-versus-host-disease model. Stem Cell Research & Therapy. 2024 Oct 29;15(1):381.

15. Mendiratta M, Mendiratta M, Rai S, Gupta R, Bakhshi S, Aggarwal M, et al. A Comparative assessment of T-Cell response of Healthy donors and Acute Graft-versus-Host-Disease Patients: Customizing immune monitoring platform [Internet]. bioRxiv; 2024 [cited 2024 Dec 24]. p. 2024.09.05.611044. Available from: https://www.biorxiv.org/content/10.1101/2024.09.05.611044v1

16. Mendiratta M, Mendiratta M, Mohanty S, Sahoo RK, Prakash H. Breaking the graft-versus-host-disease barrier: Mesenchymal stromal/stem cells as precision healers. International Reviews of Immunology. 2024 Mar 3;43(2):95–112.

17. Trigo CM, Rodrigues JS, Camões SP, Solá S, Miranda JP. Mesenchymal stem cell secretome for regenerative medicine: Where do we stand? Journal of Advanced Research [Internet]. 2024 May 9 [cited 2025 Jan 28]; Available from: https://www.sciencedirect.com/science/article/pii/S2090123224001814

18. Muzes G, Sipos F. Mesenchymal Stem Cell-Derived Secretome: A Potential Therapeutic Option for Autoimmune and Immune-Mediated Inflammatory Diseases. Cells. 2022 Jul 26;11(15):2300.

19. González-González A, García-Sánchez D, Dotta M, Rodríguez-Rey JC, Pérez-Campo FM. Mesenchymal stem cells secretome: The cornerstone of cell-free regenerative medicine. World J Stem Cells. 2020 Dec 26;12(12):1529–52.

20. Ferreira JR, Teixeira GQ, Santos SG, Barbosa MA, Almeida-Porada G, Gonçalves RM. Mesenchymal Stromal Cell Secretome: Influencing Therapeutic Potential by Cellular Pre-conditioning. Front Immunol. 2018;9:2837.

21. Teng CCJ, Kukumberg M, Rufaihah* AJ. Therapeutic Implications of Stem Cell Secretome. Journal of Stem Cell Therapy and Transplantation. 2024 Jun 17;8(1):029–32.

22. Song N, Scholtemeijer M, Shah K. Mesenchymal Stem Cell Immunomodulation: Mechanisms and Therapeutic potential. Trends Pharmacol Sci. 2020 Sep;41(9):653–64.

23. Gao F, Chiu SM, Motan D a. L, Zhang Z, Chen L, Ji HL, et al. Mesenchymal stem cells and immunomodulation: current status and future prospects. Cell Death & Disease. 2016 Jan;7(1):e2062–e2062.

24. Caplan AI, Correa D. THE MSC: AN INJURY DRUGSTORE. Cell Stem Cell. 2011 Jul 8;9(1):11–5.

25. Le Blanc K, Mougiakakos D. Multipotent mesenchymal stromal cells and the innate immune system. Nat Rev Immunol. 2012 May;12(5):383–96.

26. Arki MK, Moeinabadi-Bidgoli K, Niknam B, Mohammadi P, Hassan M, Hossein-Khannazer N, et al. Immunomodulatory performance of GMP-compliant, clinical-grade mesenchymal stromal cells from four different sources. Heliyon. 2024 Jan 30;10(2):e24948.

27. Kim DW, Staples M, Shinozuka K, Pantcheva P, Kang SD, Borlongan CV. Wharton’s Jelly-Derived Mesenchymal Stem Cells: Phenotypic Characterization and Optimizing Their Therapeutic Potential for Clinical Applications. Int J Mol Sci. 2013 May 31;14(6):11692–712.

28. Vohra M, Arora SK. Mesenchymal stem cellsthe master immunomodulators. Explor Immunol. 2023 Apr 27;3(2):104–22.

29. Han Y, Yang J, Fang J, Zhou Y, Candi E, Wang J, et al. The secretion profile of mesenchymal stem cells and potential applications in treating human diseases. Sig Transduct Target Ther. 2022 Mar 21;7(1):1–19.

30. Zhou C, Bai XY. Strategies for the induction of anti-inflammatory mesenchymal stem cells and their application in the treatment of immune-related nephropathy. Front Med [Internet]. 2022 Aug 19 [cited 2025 Jan 28];9. Available from: https://www.frontiersin.org/journals/medicine/articles/10.3389/fmed.2022.891065/full

31. Ferrara JL, Levine JE, Reddy P, Holler E. Graft-versus-host disease. The Lancet. 2009 May 2;373(9674):1550–61.

